# Achieving spatial multi-omics integration from unaligned serial sections with DIME

**DOI:** 10.64898/2026.02.25.707961

**Authors:** Pengyu Sun, Xinlei Huang, Tian Mou, Xubin Zheng

## Abstract

Learning integrated representations from spatial multi-omics data is a fundamental challenge, particularly in the context of “diagonal integration”, where data are collected from serial tissue sections across distinct omics modalities. Existing methods typically rely on the assumption of feature intersection to construct a common metric space, a prerequisite that is absent in this setting. To address this, we propose the **D**iagonal **I**ntegration Model for Spatial **M**ulti-omics **E**mbedding (DIME), a novel deep learning framework that couples a graph contrastive learning objective with cross-modal correspondence. This global correspondence is established by a hybrid alignment strategy: it first anchors high-confidence regions using Coherent Point Drift with Linear Assignment, and then extends matching to the entire tissue manifold via an Optimal Transport formulation encoding relative geodesic distances. Designed to balance inter-modal guidance with intra-modal structure preservation, DIME enables robust fusion and denoising. Experiments on simulated and real human tissue datasets demonstrate DIME’s superior robustness and versatility, where its learned representations achieve outstanding clustering accuracy and unlock the identification of biologically meaningful spatial domains. The code is available at: https://github.com/Spyyyyyyy/DIME

## 1 INTRODUCTION

Spatial omics technologies are fundamentally reshaping our understanding of the tissue microenvironment by resolving cellular heterogeneity and functional organization [1, 2]. Recent technological advancements, such as spatial CUT&Tag-RNA-seq [3] for simultaneous detecting spatial transcriptomics and epigenomics, and Stereo-CITE-seq for detecting spatial transcriptomics and proteomics. Integrating these complementary modalities aims to construct a more comprehensive omics representation to elucidate spatial gene regulation and microenvironment dynamics in complex tissues from a systems perspective [4, 5]. In response, a suite of computational methods has been developed for “horizontal” (e.g. GraphST [6], STAGATE [7], SpaGCN [8]) and “vertical” (e.g. totalVI [9], SpatialGlue [4]) integration to fuse multi-modal data and decipher anatomical details.

However, simultaneous multi-modal profiling faces significant experimental trade-offs between the “depth” and “breadth” of information [10]. A more pragmatic and powerful paradigm involves acquiring high-depth, single-modality data from serial (adjacent) tissue sections. While this strategy maximizes the biological resolution of each discrete assay, it generates spatially unaligned datasets, rendering conventional, alignment-presupposing integration methods ineffective [11]. While “mosaic” integration methods (e.g., Cobolt [12], StabMap [13], MIDAS [14], SpaMosaic [11]) have emerged, they share a critical limitation: they all require a shared modality to serve as a “bridge” for alignment. They fundamentally fail in the challenging scenario where no such bridge exists.

We define this unaddressed, bridge-free task of integrating completely disparate modalities from unaligned serial sections as “diagonal integration”. Technically, existing methods rely on the assumption of feature intersection to construct a common metric space. In the absence of such a shared feature space (the “bridge”), we encounter a dual technical challenge. First, the spatial misalignment of the serial sections fundamentally destroys the necessary morphological correspondence between samples. Second, the feature distributions are strictly disjoint (modality inconsistency), which eliminates the crucial anchor points required for feature-based alignment. This combined absence of structural and feature correspondence renders the joint optimization of robust representation learning and inter-modal integration mathematically intractable for conventional feature-anchored frameworks.

In this work, we propose DIME, a **D**iagonal **I**ntegration Model for Spatial **M**ulti-omics **E**mbedding based on Graph Neural Network (GNN) [15]. Its core premise is that the shared morphological structure of serial sections provides a powerful inductive bias, even without a common modality. This framework employs a hybrid-strategy alignment module built upon Coherent Point Drift (CPD) [16] and Optimal Transport (OT), which leverages this shared structure to estimate a cross-modal sample correspondence. Critically, DIME does not treat this correspondence as a hard label. Instead, it is employed as a regularizer within a novel graph structure contrastive loss, which robustly balances inter-modal guidance against the preservation of intra-modal neighborhood topology. We validated DIME on both simulated and real-world multi-omic datasets, integrating transcriptomic (RNA) and proteomic (Antibody Derived Tags, ADT) data. Across all benchmarks, DIME demonstrated superior performance in clustering accuracy and spatial domain identification, establishing it as the first framework to successfully address the diagonal integration challenge. Notably, DIME resolved key biological structures (e.g., T-cell zones and B-cell compartments) obscured by noise or over-smoothing in competing methods, confirming its ability to generate spatially coherent and biologically meaningful representations. Our contributions are summarized as follows:

- We propose DIME, the first framework for the diagonal integration of spatial multi-omics data from serial sections that lack a shared modality.
- We introduce a hybrid spatial alignment strategy, combining Coherent Point Drift and Optimal Transport, to estimate a robust correspondence via the construction of morphological pseudo-anchors.
- We design a novel graph structure contrastive loss that balances the cross-modal prior against intramodal structure preservation, thereby enabling effective fusion by matching neighborhood distribution.

## 2 RESULTS

### 2.1 Overall Structure of DIME

To tackle the ill-posed nature of diagonal integration, where omics data from serial sections are both spatially unaligned and feature-disparate (Fig 1), we propose DIME. The core innovation of our framework lies in strategically decoupling the dual alignment problem. Given the absence of a bridging modality, we prioritize geometric consistency to resolve the ambiguity.

**Fig. 1.**
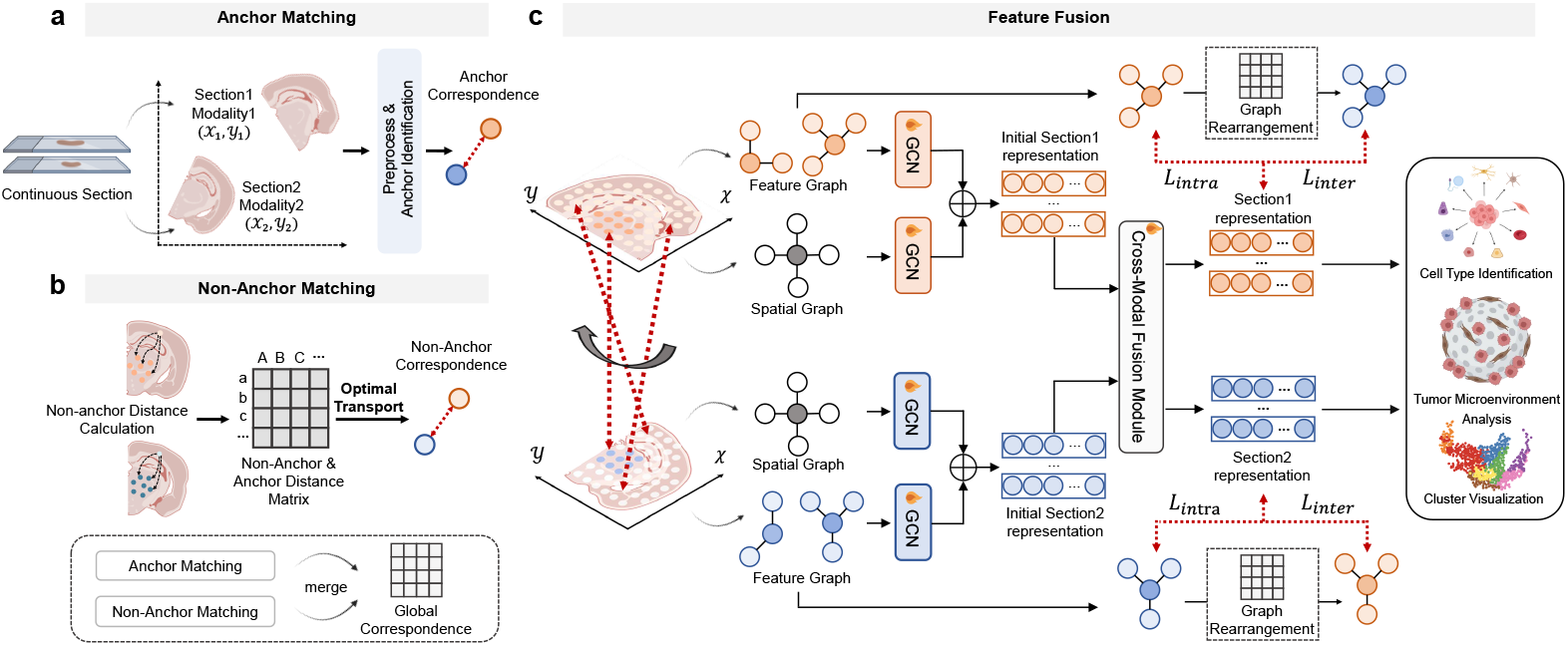
The DIME framework for diagonal integration of spatial multi-omics. (a,b) The hybrid-strategy spatial alignment module. Morphological features are used to identify and match anchor clusters. Geodesic distance and OT are then used to align non-anchor regions, yielding a global correspondence map. **(c) The prior-guided graph neural network**. A dual-graph encoder and cross-attention module fuse omics data, guided by an intra-modal structure preservation loss (ℒ_*intra*_) and an inter-modal prior-guided loss (ℒ_*inter*_).

The spatial misalignment is addressed by our hybrid spatial alignment module (Sec 4.1). We first independently cluster the omics expression profiles of the spots to generate sets of spatial clusters. Leveraging the shared morphological structure revealed by these clusters, the module integrates CPD with Linear Assignment to identify high-confidence anchors by modeling the alignment as a maximum likelihood estimation of a Gaussian Mixture Model (GMM). Subsequently, to propagate these alignments to non-anchor points, we employ an OT formulation. Crucially, unlike standard OT which relies on Euclidean metrics, we construct a structure-aware cost matrix that encodes the relative geodesic distance on the tissue manifold. Such a design provides robust global pairwise correspondence.

This preliminary correspondence then acts as a crucial inter-modal guidance signal for the subsequent GNN fusion module (Sec 4.2). Designed to process inputs from serial sections with distinct modalities, this module overcomes feature disparity using a dual-graph (feature and spatial) encoder coupled with a crossattention mechanism. Crucially, DIME effectively integrates complementary features from the alternate modality onto the respective spatial coordinates of each section, thereby bridging the information gap across the unaligned tissues. The embedding process is optimized via our graph structure contrastive loss ℒ_*intra*_ and ℒ_*inter*_, which is uniquely designed to robustly balance the inter-modal guidance from the spatial alignment against the essential preservation of intrinsic intra-modal neighborhood topology. Ultimately, DIME generates a unified, spatially coherent latent representation that significantly enhances performance in downstream tasks, such as cell type identification.

### 2.2 DIME Robustly Integrates Simulated Unaligned Sections

We first validated DIME on a simulated dataset designed with complementary, unaligned RNA and ADT patterns to test diagonal integration. Qualitative analysis (Fig 2a) shows that while baseline methods (e.g., Principal Components) fail to delineate all spatial regions, DIME successfully reconstructs the complete ground-truth pattern. By leveraging the sample correspondences derived from its spatial alignment framework, DIME effectively imputes the missing factors in each section, a task where general integration frameworks fail.

**Fig. 2.**
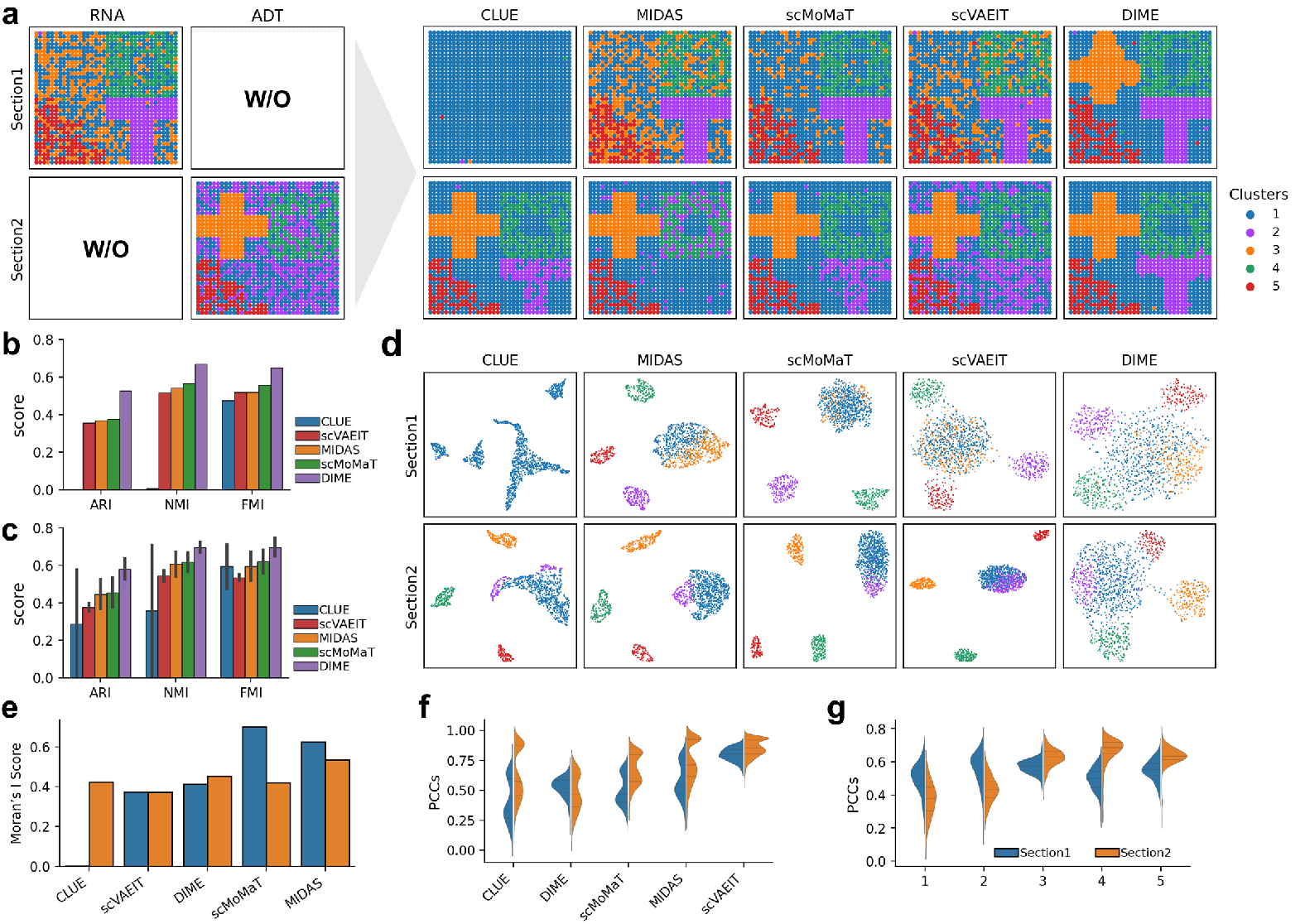
DIME enables effective integration and denoising of simulated diagonal multi-omics datasets. (a) Spatial clustering results of baseline methods and DIME on the diagonal dataset. Results are shown for Section 1 (RNA-only) and Section 2 (ADT-only), highlighting DIME’s ability to recover structural features via cross-modal imputation. (b,c) Quantitative evaluation of clustering performance (ARI, NMI, and FMI) using Section 1 (b) and Section 2 (c) as the spatial template for integration, respectively. (d) UMAP visualization of integrated latent representations for Section 1 (top row) and Section 2 (bottom row), demonstrating superior cluster separation by DIME. (e) Evaluation of spatial coherence across integrated sections using Morans *I* statistic. (f,g) Pearson correlation coefficient (PCC) analysis of embedding distance matrices, showing the overall correlation performance across methods (f) and the cluster-specific correlation results (g).

We then quantitatively evaluated clustering accuracy using the Adjusted Rand Index (ARI), Normalized Mutual Information (NMI), and Fowlkes-Mallows Index (FMI). Compared to using the original expression patterns, DIME’s integrated embeddings show a marked improvement in all metrics and substantially outperform competing methods (Fig 2b, c).

Finally, we assessed the spatial coherence and feature preservation of the embeddings. Notably, while certain baseline methods achieved comparable or even higher Moran’s I scores and Pearson Correlation Coefficients (PCC) in specific regions (Fig. 2e, f), their qualitative spatial clustering (Fig. 2a) revealed a profound inability to reconstruct the correct biological topology. This discrepancy highlights the limitations of global metrics like Moran’s I and PCC in the context of diagonal integration; a high Moran’s I can be an artifact of excessive spatial smoothing that obscures fine-grained boundaries, whereas a high PCC may merely reflect the over-fitting of noisy, original features (as seen with scVAEIT) rather than the successful recovery of cross-modal information. In contrast, DIME strikes a superior balance by constructing spatially-aware embeddings that prioritize the accurate restoration of the tissue’s geometric and biological “skeleton,” thereby ensuring both global coherence and local fidelity.

### 2.3 DIME Integrates Diagonal Sections to Reconstruct the Human Lymph Node

To rigorously evaluate DIME’s performance in a complex, non-aligned context, we constructed an unpaired spatial multi-omic dataset from adjacent human lymph node sections, integrating RNA (Section 1) and ADT (Section 2). Spatial visualization of the integrated results (*k* = 7 clusters) highlights DIME’s superior ability to resolve intricate tissue architectures compared to existing methods (Fig 3a). Baseline models exhibited significant limitations: CLUE suffered from pronounced over-smoothing in Section 1 and failed to identify the medulla-related region (clusters 1, 2); MIDAS retained substantial noise across both sections; and scMoMaT showed inconsistent performance. While scVAEIT identified some broad tissue regions, DIME provided higher-fidelity delineation, uniquely resolving the cortex (cluster 5) in Section 1 and more precisely capturing the capsule (cluster 3). This qualitative superiority is mirrored in the latent space, where DIME’s UMAP embedding (Fig 3c) shows well-defined cluster boundaries that strongly correspond to ground-truth histological labels.

**Fig. 3.**
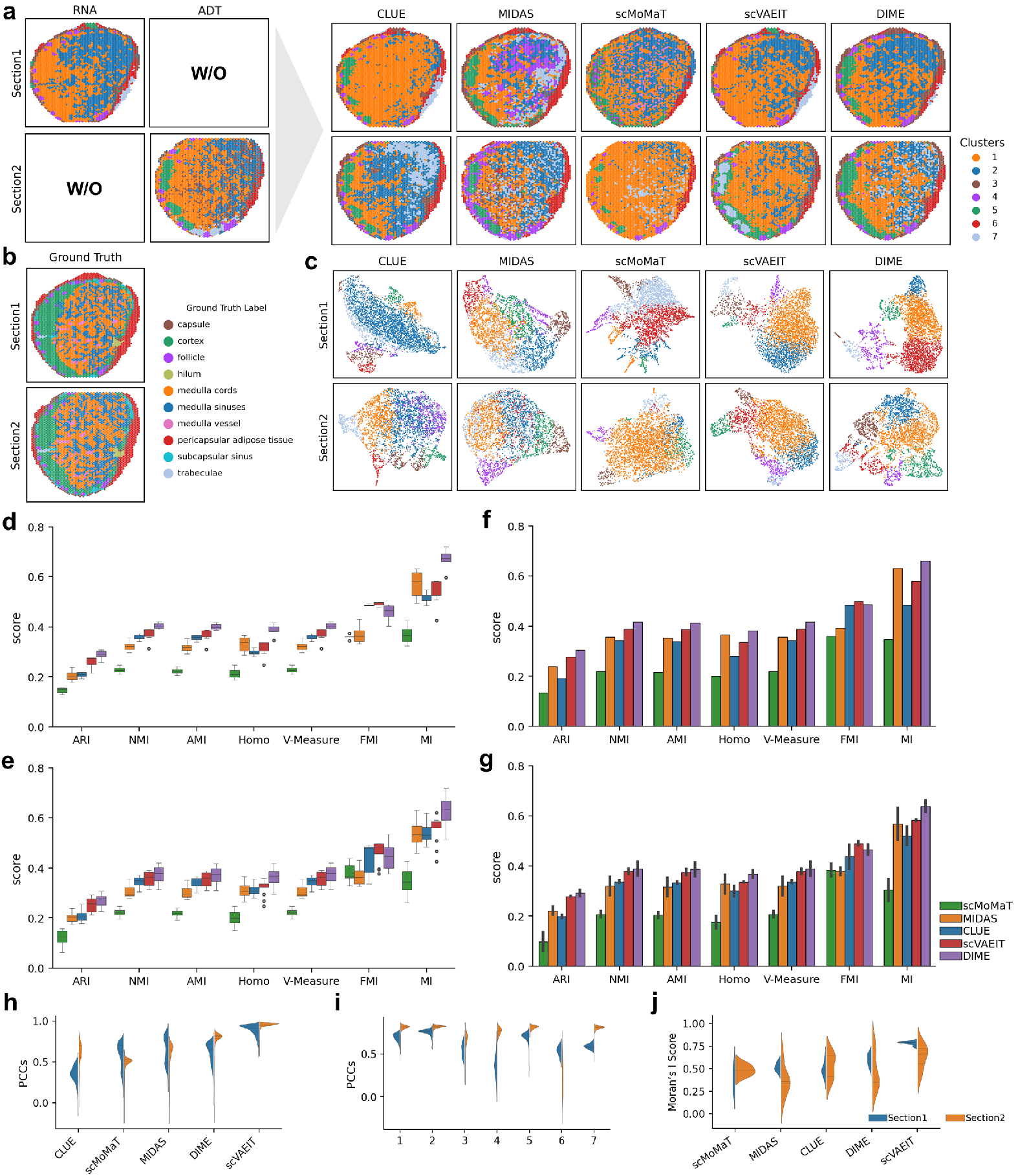
Integration and quantitative evaluation on the human lymph node dataset. (a) Spatial clustering results (at *k* = 7) comparing single-modality profiles, baseline integration methods, and DIME. (b) UMAP visualization of the integrated latent representations, showing the blending of modalities and cluster distinctness. (c) Ground truth manual annotations provided for the lymph node sections. (d,e) Robustness analysis of clustering performance (ARI and NMI) across varying resolutions (*k* = 4 to 10) using Section 1 (d) and Section 2 (e) as the integration template. (f,g) Quantitative comparison of clustering metrics at the optimal resolution (*k* = 7) for Section 1 (f) and Section 2 (g). (h,i) Pearson correlation coefficient (PCC) analysis of embedding distance matrices, depicting overall method performance (h) and cluster-specific correlation (i). (j) Evaluation of spatial coherence for the integrated results using Morans *I* statistic.

Beyond this visual inspection, we assessed model stability and quantitative performance. DIME consistently outperformed all baseline methods across a range of clustering resolutions (*k* = 4 to 10), underscoring its robustness (Fig 3d, e). At the *k* = 7 resolution, DIME demonstrated a clear advantage across standard clustering evaluation metrics (Fig 3f, g). We further assessed spatial coherence using Moran’s I (Fig 3j). DIME achieved the second-highest score; notably, the top-scoring method (scVAEIT) produced spatially “clean” but histologically incorrect partitions (as seen in Fig 3b). This suggests its high Moran’s I score is an artifact of over-smoothing, whereas DIME’s strong performance represents a more optimal balance between capturing true spatial patterns and preserving high-fidelity, complex tissue boundaries. Finally, to interpret the integrated latent space, we computed PCCs between the original modalities and the DIME-generated embeddings (Fig 3h, i), revealing that protein features provided a stronger contribution to the final representation in most tissue-specific clusters.

### 2.4 Biological Validation of DIME’s Integration in Human Tonsil

To further validate DIME, we integrated unpaired spatial RNA-seq (Section 1) and proteomics (Section 2) data from human tonsil tissue. Visual analysis of the six identified clusters (*k* = 6) immediately revealed significant performance disparities (Fig 4a). Integration results from MIDAS and scMoMaT were dominated by spatial noise, rendering them unusable. Conversely, CLUE and scVAEIT suffered from severe over-smoothing, which obliterated key histological features and led to their failure in delineating Connective tissue (cluster 5) in Section 1 and the Epithelial region (cluster 3) in Section 2. In sharp contrast, DIME achieved an optimal balance of noise reduction and feature preservation. In Section 1, it precisely resolved the Follicle mantle zone (cluster 4) and accurately delineated both Connective tissue (cluster 5) and Epithelial (cluster 3) regions. In Section 2, it faithfully reconstructed the protein-defined tissue structures, including the corresponding Connective tissue (cluster 5). This qualitative superiority was confirmed by quantitative metrics, where DIME achieved the optimal performance, significantly outperforming all other methods (Fig 4b, c).

**Fig. 4.**
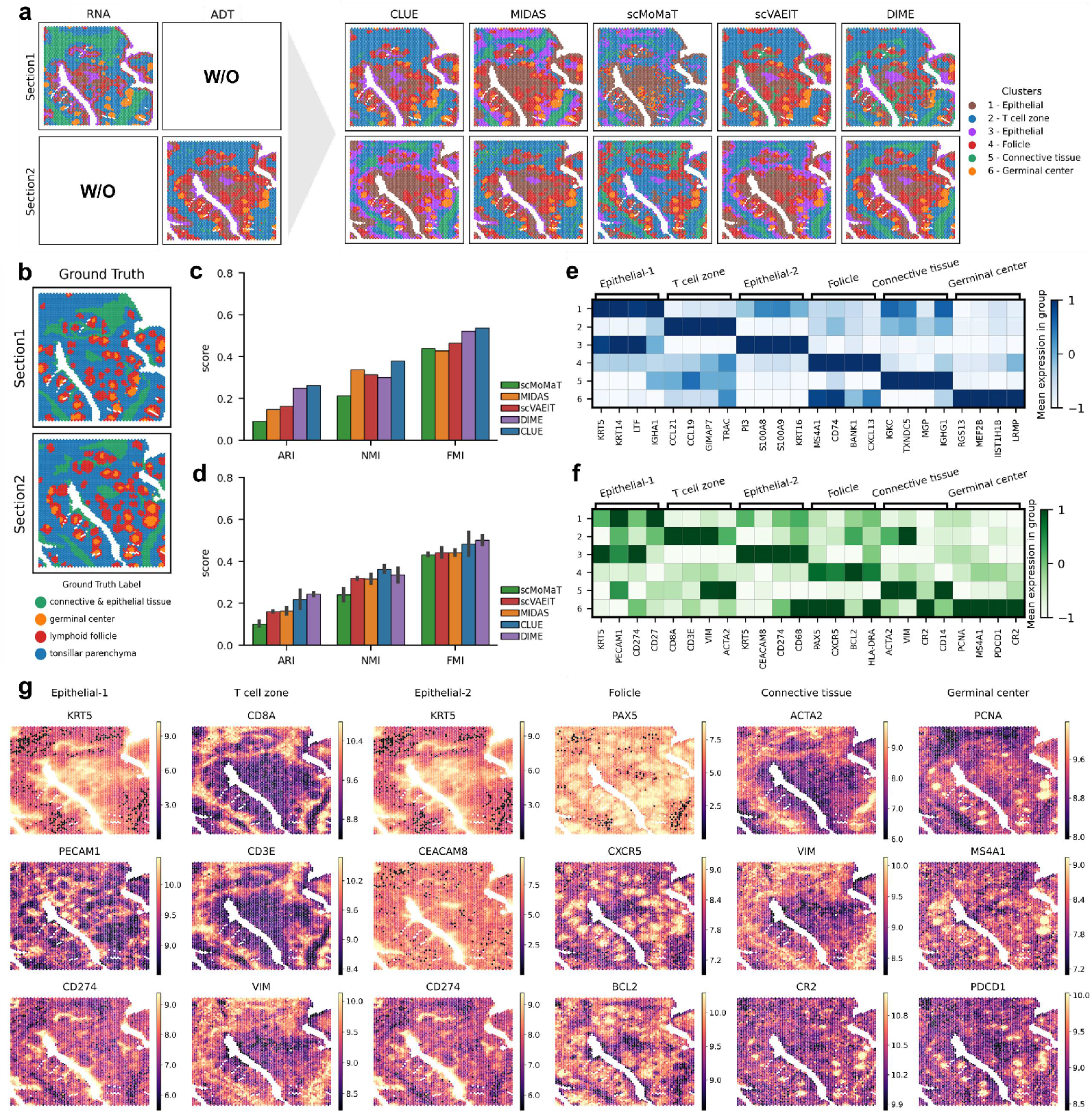
Integration and biological validation on the human tonsil dataset. (a) Spatial clustering results (at *k* = 6) comparing single-modality profiles, baseline methods, and DIME. (b) Ground truth manual annotations of the tonsil tissue sections. (c,d) Quantitative evaluation of clustering performance (ARI, NMI, and FMI) using Section 1 (c) and Section 2 (d) as the spatial integration template. (e,f) Biological validation of DIME-identified clusters through expression heatmaps of region-specific RNA markers (e) and ADT surface markers (f), confirming the accurate identification of distinct biological compartments (e.g., germinal centers and T-cell zones). (g) Visualization of normalized ADT levels for key lineage-specific surface markers across identified spatial domains.

To confirm the biological relevance of DIME’s integrated clusters, we analyzed canonical marker gene and protein expression (Fig 4e, f, g). The results showed strong biological fidelity: the “T cell zone” cluster correctly co-expressed T-cell RNAs (e.g., TRAC) and proteins (e.g., CD3E/CD8A). Furthermore, DIME successfully deconstructed B-cell compartments, identifying the “Follicle” (defined by the CXCL13-CXCR5 axis) and the “Germinal center” (containing proliferating B-cells and T-cells) as distinct multicellular units. This analysis confirms DIME’s ability to generate not only spatially coherent but also biologically meaningful and interpretable representations from unpaired multi-omic data.

## 3 DISCUSSION

In this study, we presented DIME, a deep learning framework designed to address the formidable challenge of “diagonal integration” in spatial multi-omics. By decoupling spatial registration from molecular feature matching, DIME overcomes the fundamental limitation of existing methods that rely on shared feature spaces. The core of our approach lies in a hybrid-strategy spatial alignment module that leverages the physical manifold of tissue sections, ensuring robust cross-modal correspondence even when biological signatures exhibit significant heterogeneity.

A key strength of the DIME architecture is its inherent flexibility in handling diverse omics granularities. Unlike traditional methods that assume a one-to-one correspondence between sequencing spots or require identical clustering resolutions, DIME allows for independent cluster numbers (*n*_*S*_ ≠ *n*_*T*_) across modalities. This asymmetric matching capability is particularly crucial when integrating technologies with disparate feature densities, such as highly multiplexed RNA-seq and relatively sparse ADT profiles. By prioritizing the alignment of stable histological macro-structuresidentified through morphological descriptors and relative spatial positioningDIME ensures that the integration is grounded in the underlying tissue architecture rather than being susceptible to local molecular noise or clustering stochasticity.

Furthermore, the DIME framework is fundamentally modality-agnostic, offering significant potential for expansion beyond dual-modal integration. The hybrid alignment modulecombining morphological landmarks with geodesic-based Optimal Transportoperates on the geometric manifold of the tissue. Consequently, the current pairwise alignment of RNA and ADT can be readily generalized to a tri-modal or multi-modal diagonal integration framework. For instance, DIME could simultaneously bridge RNA, ADT, and ATAC-seq data from consecutive sections by incorporating specialized encoding branches tailored to each modalitys unique feature profile. This extensibility positions DIME as a versatile backbone for constructing comprehensive, high-definition spatial atlases that capture multiple layers of biological regulation

However, certain areas warrant further investigation to fully realize the model’s potential. While DIME demonstrates the theoretical and practical feasibility of multi-modal diagonal integration, future work will focus on optimizing the computational efficiency of geodesic distance calculations as spatial omics datasets scale to millions of spots. Additionally, exploring the direct application of DIME to non-consecutive sections or tissues with significant mechanical deformation will further broaden its utility in complex pathological studies.

In conclusion, DIME provides a robust and extensible solution for the diagonal integration of spatial multi-omics data. By bridging the gap between disparate molecular spaces through spatial-topological priors, DIME facilitates a more comprehensive understanding of the intricate regulatory processes within the spatial context of complex biological systems.

## 4 METHODS

### 4.1 Hybrid Spatial Alignment for Structural Prior Estimation

Recognizing that a perfect alignment is ill-posed under non-rigid deformations, we aim to estimate high-confidence spatial correspondences to serve as an inductive bias for the fusion network. Crucially, direct alignment based on raw spatial coordinates is often inadequate due to inevitable mechanical artifactssuch as stretching, shearing, and micro-scale distortionsintroduced during sectioning and mounting. Unlike conventional rigid registration, which assumes uniform transformations and is sensitive to these distortions, DIME prioritizes topological continuity over absolute geometric proximity. By leveraging morphological descriptors and relative positioning, DIME establishes a feature-aware coordinate system that remains resilient to physical deformations, ensuring that cross-modal correspondences are grounded in true biological homology.

#### Anchor Cluster Identification

Let *S* and *T* represent the sets of spatially resolved spots from two serial sections characterized by different omics modalities, designated as the source and target, respectively. To bridge these distinct molecular spaces, we first independently partition the spots into *n* spatial domains, 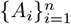 for *S* and 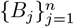 for *T*, based on their respective omics expression profiles. Although cross-modal biological signatures may exhibit heterogeneity, the macroscopic histological architecture remains physically coupled across serial sections. Leveraging this structural continuity, we extract a comprehensive set of morphological descriptors for each domainincluding area, eccentricity, perimeter-to-area ratio, and principal axis orientationderived directly from the spatial coordinates of the constituent spots. To further ensure the reliability of the correspondence, we incorporate the relative spatial positioning of these domains as a global topological constraint. By minimizing the Euclidean distance within this joint morphological and positional feature space, we identify the optimal pair of anchor clusters (*A, B*) that exhibits the highest structural homology. This strategy effectively utilizes the stable geometric “skeleton” of the tissue to provide a robust, coarse-to-fine initialization, ensuring that the subsequent sparse-to-dense spatial matching remains resilient to local molecular noise or clustering inconsistencies.

#### Anchor Matching via Coherent Point Drift and Linear Assignment

To establish robust point-level correspondence within the identified anchor clusters, we employ a two-stage strategy: probabilistic registration followed by combinatorial optimization. Let the source anchor cluster be 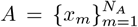 and the target anchor cluster be 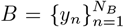, where *N*_*A*_ and *N*_*B*_ denote the number of spots in the respective clusters.

##### 1. Probabilistic Non-rigid Registration via Coherent Point Drift

We formulate the alignment of the source set 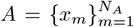 and target set 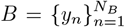 (both embedded in ℝ^*D*^, where *D* = 2 represents the 2D spatial coordinates of the spots) as a probability density estimation problem, adopting the framework of CPD [16]. The core idea is to treat the transformed source points 𝒯 (*x*_*m*_, *V*) = *x*_*m*_ + *v*(*x*_*m*_) as centroids of a GMM that generates the target set *B*. The non-rigid displacement field *v* is parameterized by the coefficient matrix 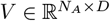.

The registration likelihood is modeled by the Probability Density Function (PDF) *p*(*y*) for observing a target point *y*. Following [16], the PDF is defined as a mixture of *N*_*A*_ Gaussian components (inliers) and one uniform outlier component:

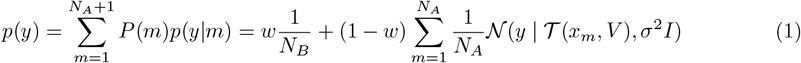

where *w* is the outlier weight and *σ*^2^ is the isotropic variance. The probability density function *p*(*y*) is a GMM, whose expectation is calculated by the summation 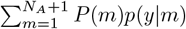. This term explicitly incorporates the outlier and inlier components, using the respective priors *P* (*m*) defined as follows:

- **Inliers (***m* ≤ *N*_*A*_): The prior is 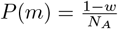, and the likelihood is *p*(*y*|*m*) = 𝒩 (*y*|𝒯 (*x*_*m*_, *V*), *σ*^2^*I*).
- **Outlier (***m* = *N*_*A*_ + 1): The prior is *P* (*m*) = *w*, and the likelihood is 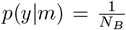 (uniform density).

Based on this probabilistic model, we estimate the parameters *θ* = {*V, σ*^2^} by minimizing the regularized negative log-likelihood:

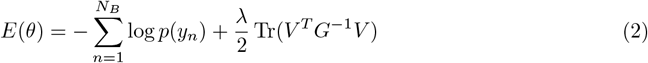

The first term is the Negative Log-Likelihood, where *p*(*y*_*n*_) is the PDF evaluated at the *n*-th target point. The second term is the Motion Coherence constraint, governed by the Radial Basis Function kernel matrix *G*, defined as 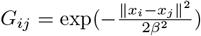. Here, *β* is the kernel bandwidth, a hyperparameter controlling the degree of spatial smoothness. This term explicitly controls the local correlation of deformations to enforce spatial smoothness. The parameter *λ* is the regularization weight, which governs the trade-off between maximizing data fidelity and minimizing deformation energy.

We optimize Eq. (2) using the Expectation-Maximization algorithm. In the **E-step**, the posterior correspondence probability *P* (*m*|*y*_*n*_) is computed:

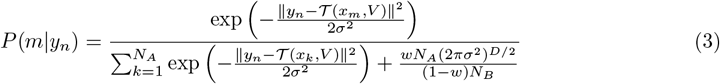

The denominator includes an explicit outlier term that acts as a robust mechanism to suppress correspondence probability for distant target points.

In the subsequent **M-step**, the parameters *θ* = {*V, σ*^2^} are updated by maximizing the expected log-likelihood. The coefficient matrix *V* is updated by solving a weighted linear system using the correspondences *P* (*m* | *y*_*n*_), and *σ*^2^ is calculated via the weighted mean squared error. Throughout the optimization, the variance *σ*^2^ is iteratively reduced, facilitating a deterministic annealing process that ensures convergence from a robust coarse alignment to fine details.

##### 2. Discrete Correspondence via Linear Assignment Problem

The posterior matrix 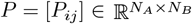 derived from the CPD framework provides soft, probabilistic correspondences, where *P*_*ij*_ represents the probability of a match between spot *i* in the source section and spot *j* in the target section. To establish unique, one-to-one discrete mappings and effectively filter out spurious matches, we formulate this task as a maximum weight linear assignment problem. The optimal binary assignment matrix *M* ^∗^ is determined by maximizing the total correspondence reward:

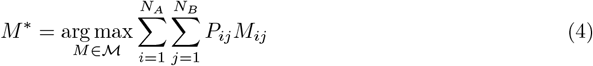

where the feasible set ℳ denotes the space of partial permutation matrices that satisfy strict one-to-one constraints:

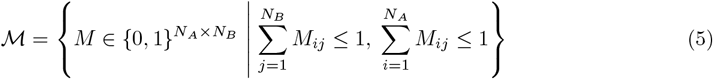

By framing the objective as a maximization of the posterior evidence, we ensure that the algorithm identifies the most confident spatial correspondences while the inequality constraints inherently accommodate unmatched points (outliers). This problem is solved efficiently using the Jonker-Volgenant (LAPJV) algorithm [17]. Finally, the definitive matched indices 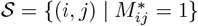 are extracted to finalize the anchor pairs (*I*_*A*_, *I*_*B*_).

#### Non-Anchor Matching via Geodesic Optimal Transport

For the remaining non-anchor regions (*S*^*′*^ = *S \ I*_*A*_ and *T* ^*′*^ = *T \ I*_*B*_), we employ an OT strategy grounded in global spatial structure. This approach relies on a novel point representation derived from the spatial manifold, which we construct in two steps:

##### 1. Spatial Graph Construction

We first model the spatial manifold of each section by constructing a *k*-nearest neighbor (*k*-NN) graph *G* = (*V, E*) from the normalized spatial coordinates. In this graph, the vertex set *V* corresponds to the *N* spatial locations (e.g., spots), and the edge set *E* captures the local connectivity of the manifold.

##### 2. Geodesic Distance Representation [18]

We approximate the geodesic distance between any two points by the shortest-path distance *d* (·, ·) on *G*. For each non-anchor point *s*_*i*_ ∈ *S*^*′*^, we define its global position relative to the source anchors *a*_*j*_ ∈ *I*_*A*_ as the vector:

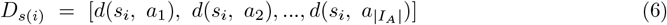

Analogously, we compute *D*_*t*(*i*)_ for every *t*_*i*_ ∈ *T* ^*′*^ relative to the target anchor points *I*_*B*_. These vectors serve as high-dimensional descriptors that encode the global position of non-anchor points relative to the stable anchor framework. Critically, the LAP (Sec 4.1) ensures a one-to-one mapping between the matched anchor points, which we denote as *I*_*A*_ ⊂ *A* and *I*_*B*_ ⊂ *B*, and guarantees that the number of matched anchors | *I*_*A*_ | = | *I*_*B*_ |. This dimensional equivalence validates the direct pairwise comparison between *D*_*s*(*i*)_ and *D*_*t*(*j*)_, forming the basis for the Euclidean distance cost matrix *C*.

##### 3. Optimal Transport

We compute the pairwise Euclidean distance between the source and target descriptors as the cost matrix *C*, specifically *C*_*ij*_ = ∥ *D*_*s*(*i*)_ − *D*_*t*(*j*)_ ∥_2_. Standard optimal transport suffers from high computational complexity. To ensure computational efficiency and differentiability, we adopt the entropy-regularized optimal transport formulation [19]. We seek the transport plan *P* ^∗^ that minimizes the total cost regularized by the entropy of the plan:

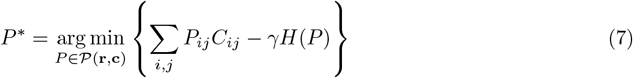

where *γ* is the regularization coefficient and *H*(*P*) = −∑_*i,j*_ *P*_*ij*_ log *P*_*ij*_ is the entropy of the transport plan. The set of feasible transport plans 𝒫 (**r, c**) is defined by the mass conservation constraints:

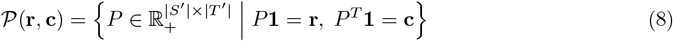

Here, **1** denotes a vector of ones (with appropriate dimension), and **r** ∈ ℝ^|*S′*|^ and **c** ∈ ℝ^|*T′*|^ are the normalized marginal distribution vectors for the non-anchor points, typically uniform 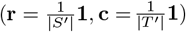. This is solved efficiently using the Sinkhorn algorithm, yielding the soft correspondence matrix *P* ^∗^ for the non-anchor regions.

#### Constructing the Global Correspondence

Finally, we construct a global, hybrid correspondence 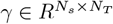:

1. **Anchor Regions:** For anchor pairs (*i, j*) identified in (*I*_*A*_, *I*_*B*_), we set *γ*_*ij*_ = 1 and *γ*_*ik*_ = 0 for all *j* ≠ *k*, enforcing a hard one-to-one mapping.

2. **Non-Anchor Regions:** For *i* ∈ *S*^*′*^ and *j* ∈ *T* ^*′*^, we set *γ*_*ij*_ = (*P* ^∗^)_*i*_*′*_*j*_*′*, where *i*^*′*^ and *j*^*′*^ are the corresponding indices in the non-anchor sets *S*^*′*^ and *T* ^*′*^.

From *γ*, we derive a final single-valued (hard) mapping *f* : *S* → *T*. For *i* ∈ *I*_*A*_, *f* (*i*) is uniquely determined. For *i* ∈ *S*^*′*^, we use Maximum Probability Assignment:

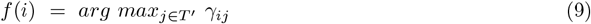

This yields a complete deterministic correspondence map *f*, which guides the spatial graph rearrangement during the fusion network’s loss computation.

### 4.2 Prior-Guided Graph Neural Network for Fusion

We propose a GNN-based fusion network, architecturally composed of an intra-modal encoder and a cross-modal fusion module. This network is optimized by our novel Guiding Graph Structure Loss. This loss function is explicitly designed to balance two competing objectives: (1) an Intra-modal Structure Preservation term (ℒ_*intra*_) and (2) an Inter-modal Information Fusion term (ℒ_*inter*_). Crucially, this ℒ_*inter*_ term is guided by the correspondence map *f* to enforce cross-modal consistency.

#### Intra-modal Dual-Graph Encoder

For each modality *m*, we begin by constructing two distinct adjacency matrices. First, we define an omic-specific feature adjacency matrix 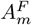 based on feature similarity. Second, we construct a spatial adjacency matrix 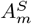 by applying the *k*-NN algorithm to the physical coordinates of the spots.

We then employ two parallel Graph Convolutional Network (GCN) [15] encoders to generate specialized embeddings from the omic feature matrix *X*_*m*_, using each adjacency matrix respectively:

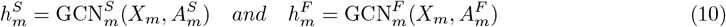

Here, 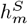 captures the spatially-aware representation, while 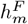 captures the feature-centric representation.

Inspired by SpatialGlue [4], we fuse these two representations using an intra-modal attention mechanism. This layer computes scalar attention weights, 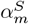 and 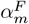, which learn to balance the relative contributions of the spatial and feature-based embeddings for each modality. The initial latent representation, 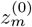, is then obtained as their weighted sum:

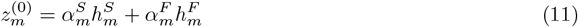

#### Cross-Modal Fusion Module

Given the initial representations 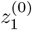 and 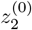 for modalities *m*_1_ and *m*_2_, we use a cross-attention module to fuse them. For *m*_1_, the update is:

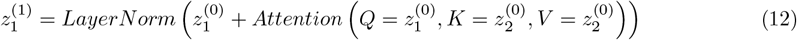

where *Q, K, V* are standard inputs to a multi-head attention block. The output 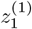 is passed through an Feed-Forward network with a residual connection to obtain the final representation *Z*_1_. A symmetric operation yields *Z*_2_.

#### Prior-Guided Training Objective

The network is trained by aligning the latent similarity distribution with the neighborhood distribution from the original feature graphs.

**1. Target Distribution (***W* ^*′*^): For a feature graph *G*^*F*^ with weighted adjacency matrix *W*, we define the target neighborhood distribution *W* ^*′*^_*ij*_ = *W*_*ij*_*/* ∑_*k*_ *W*_*ik*_ as its row-wise normalization.

**2. Latent Distribution (***P* ): For a latent representation *Z* (with rows *z*_*i*_), we compute the latent similarity distribution *P* (with elements *p*_*ij*_) using a temperature-scaled (*τ*) dot product:

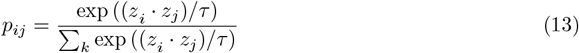

**3. Loss Function:** We minimize the difference between *W* ^*′*^ and *P* using the cross-entropy loss:

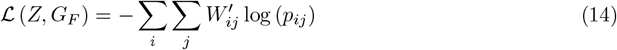

#### Total Loss

The final objective ℒ_*total*_ is a weighted sum of two loss components for each modality. For *m*_1_:

- **Intra-modal Structure Preservation Loss ((**ℒ_*inter*,1_)): This loss anchors the latent representation to its own feature graph 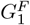. It is computed as 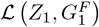, preserving modality-specific information.
- **Inter-modal Information Fusion Loss (**ℒ_*inter*,1_): This loss integrates *m*_2_’s structure into *m*_1_ using the correspondence 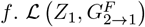 uses the correspondence map *f* : *S* → *T* (from Sec 4.1) to translate the neighborhood structure from *m*_2_ onto *m*_1._ The target distribution 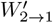 is constructed such that the desired similarity between points *i* and *j* in *m*_1_ is set to the actual neighborhood similarity of their mapped counterparts, *f* (*i*) and *f* (*j*), in *m*_2_. Formally: 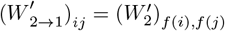.

Symmetric losses ℒ_*intra*,2_ and ℒ_*inter*,2_ are computed for *m*_2_. The total loss is:

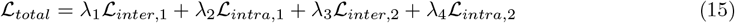

where *λ*_1_, …, *λ*_4_ are hyperparameters. This loss architecture is designed to promote denoising. We assume that modality-specific noise in 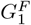 (captured by ℒ_*intra*,1_) is uncorrelated with the noise in 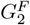 (ℒ_*inter*,1_). By balancing these two objectives, the network is trained to find a consensus representation that isolates the shared biological signal while attenuating the uncorrelated, modality-specific noise.

### 4.3 Experimental Design

#### Datasets

We conducted experiments on three spatial multi-omics datasets: a synthetic benchmark with known ground truth, and two real-world tissue datasets (Human Lymph Node and Tonsil) featuring complex biological architectures:

##### Simulated Dataset

Multivariate count data were simulated following [20] to replicate the “ggblocks” spatial pattern on a 36 × 36 grid (*N* = 1296). We simulated two modalities: RNA (*J*_RNA_ = 200) and ADT (*J*_ADT_ = 100). RNA counts follow a Zero-Inflated Negative Binomial (ZINB) distribution, while ADT counts follow a Negative Binomial (NB) distribution. Both share a mean structure **M** = **b** + **SFW**, where **F** defines factor activation, **W** denotes feature weights (10 for RNA, 12 for ADT), and **S** governs spatial factor mixing. Shape parameters were set to 20 (RNA) and 10 (ADT), with an additional dropout *p*_RNA_ for the RNA modality.

##### Human Dataset

These data sets were generated using the 10x Genomics CytAssist Visium platform, a cutting-edge technology facilitating the simultaneous spatial profiling of the transcriptome (18, 085 genes) and multiplexed protein expression. The platform utilizes next-generation sequencing (NGS) and offers a spatial resolution of 55 *µ*m with a spot spacing of 100 *µ*m. Specifically, the data employed for spatial integration consists of two human lymph node sections, A1 (3, 484 spots) and D1 (3, 359 spots), both profiling 31 proteins. The two-section **Human Tonsil** dataset, utilized for multi-section integration, includes 4, 326 and 4, 519 spots, respectively, and consistently profiles 18, 085 genes and 31 proteins across both sections. The **Human Lymph Node** dataset is publicly available at https://zenodo.org/records/17660964

#### Baseline Comparisons

Since standard spatial vertical integration methods are incompatible with spatially unaligned serial sections, we compared DIME against four State-of-the-Art (SOTA) single-cell integration methods: CLUE [21], MIDAS [22], scMoMaT [23], and scVAEIT [24]. These baselines were selected for their ability to integrate data without shared features, unlike spatial mosaic methods (e.g., SpaMosaic) which strictly rely on a bridging modality. All baseline methods were implemented using default hyperparameters. During the evaluation, the number of clusters was standardized across all methods to ensure comparability of the clustering results.

#### Evaluation Metrics

To comprehensively evaluate model performance, we selected nine metrics: Mutual Information (MI), Normalized Mutual Information (NMI), Adjusted Mutual Information (AMI), Fowlkes-Mallows Index (FMI), Adjusted Rand Index (ARI), V-measure, Homogeneity (Homo), Moran’s I, and Pearson Correlation Coefficients (PCCs) (Details in Supplementary).

## 5 ACKOWLEDGEMENTS

This work was supported by National Natural Science Foundation of China (32300554). The computational resources are supported by SongShan Lake HPC Center (SSL-HPC) in Great Bay University.

